# The Motor Wisdom of the Crowd

**DOI:** 10.1101/2022.02.04.479103

**Authors:** Gabriel Madirolas, Regina Zaghi-Lara, Adam Matic, Alex Gomez-Marin, Alfonso Pérez-Escudero

## Abstract

Wisdom of the Crowd is the aggregation of many individual estimates to obtain a better collective one. This effect has an enormous potential from the social point of view, as it means that a decision may be taken more effectively by vote among a large crowd than by a small minority of experts. Wisdom of the Crowd has been demonstrated in a wide range of cognitive tasks, all of which involve rational thinking. Here we tested this effect in the context of drawing simple geometrical shapes which, while still enacting cognitive processes, mainly involved visuo-motor control. We asked more than 700 school students to trace with their finger a predefined pattern shown on a touchscreen, and analyzed whether their individual trajectories could be aggregated in a way that improved the match with the original pattern. We found that this task has all the characteristics of the strongest examples of Wisdom of the Crowd: First, the aggregate trajectory can be up to 5 times more accurate than the individual ones. Second, this great improvement requires aggregating trajectories from different individuals (rather than different trials from the same individual). Third, the aggregate trajectory outperforms >99% of the individual trajectories. Fourth, when we split our dataset between young children (<10.5 years old) and older children, we find that older individuals outperform younger ones, as naively expected. However, a crowd of young children outperforms the average older individual. In sum, we demonstrate for the first time the Wisdom of the Crowd phenomenon in the realm of motor control, opening the door to further studies of human but also animal behavioral trajectories and their mechanistic underpinnings.

**Significance statement:** Wisdom of the Crowd is the aggregation of many individual estimates to obtain a better collective one. Thanks to a combination of mathematical, psychological and social factors, this collective estimate can be surprisingly accurate, even when it comes from a crowd of poorly informed individuals. Originally proposed in the context of simple tasks (such as estimating a number), a general version of this concept now pervades our society, from being a key reason behind the success of democracies to being explicitly used to generate collective high-quality knowledge, from Wikipedia to Stack Exchange. Despite this enormous practical success, academic research lags behind, with most studies focusing on simple and narrow tasks. Here we apply the concept for the first time to a sensory-motor task, testing whether the combination of many drawings traced by different subjects resembles the original template. This type of task is wildly different from previous studies of Wisdom of the Crowd, as it depends on continuous sensory-motor control, rather than explicit reasoning or discrete decision-making. We demonstrate that the effect is strong: The aggregate trajectories are more accurate than an overwhelming majority of the individual ones, and a crowd of young children outperforms the average teenager with more mature motor skills.

## Introduction

The Wisdom of the Crowd (WOC) is the notion that the aggregate opinion of a diverse group of people may be more reliable than that of an expert. This idea gained great standing in the academic community in 1906, when Sir Francis Galton showed that the average of hundreds of individual estimates about the weight of an ox matched its actual weight within 1%--far more accurately than a highly skilled farmer (1). The potential of this effect was rapidly appreciated, but applications were limited for a long time, due to the practical difficulty of gathering and aggregating individual opinions. The rapid development of communication technologies since the 1990 ‘s has removed that barrier, unleashing the potential of the WOC (2), which is now replacing individual experts in several realms, from collectively written Wikipedia articles that replace classical encyclopedia articles written by experts (3) to Stack Exchange posts written and voted by users that replace traditional manuals (4). These examples show the transformative potential of the WOC.

However, to properly grasp the non-trivial scope of the WOC and to ensure its validity, it is critical to understand the effect deeply. In particular, two key questions must be answered. The first key question is how to extract a WOC estimate from a group of people. In the traditional paradigm, subjects were as diverse as possible (5), each subject made an independent estimate (they did not communicate with each other) (6), and opinions were aggregated by averaging all individual guesses (2). The condition of independence is traditionally regarded as crucial to guarantee that systematic individual errors cancel out when aggregating several individuals (2, 6). An example of the negative effect of broken independence arises when subjects are informed of the guesses of others, and are then allowed to emit their guess or to reconsider their previous one (7). However, quantifying social influence over each individual allows to find new aggregation measures that counteract this effect (8), and to take advantage of it to improve upon the crowd estimate (9). Moreover, it has been shown how the condition of independence can be relaxed (10), and how allowing subjects to discuss before arriving to a consensus estimate can lead to improvements both at the group and individual level (11, 12). Others have argued for maximal differences (negative correlations) between subjects as the essential requisite for the WOC (13), and therefore the detection of correlations is presented as a powerful tool to improve the collective estimate when it deviates from the true value (14). Finally, methods to find subgroups of individuals whose aggregated estimate may be better than the aggregated estimate of the whole crowd have been proposed, for example based on identifying expertise within the crowd (15).

The second key question is what tasks can be performed more efficiently by a collective than by an individual. Classical demonstrations of the WOC consisted of guessing a number or choosing over a set of discrete alternatives (2, 6). While a big part of research still follows this paradigm, many studies have successfully applied WOC to more complex tasks, such estimation of multi-dimensional quantities (16), collective guessing of a sentence (17), collective edition (3), forecasting in prediction markets and prediction polls (18, 19), correct tempo of a classical music piece (20), medical diagnosis (21, 22), or drug prescription (23). However, all these studies share a common characteristic: They focus on explicitly rational tasks, in which individuals need to make a conscious estimate.

Here we investigated whether WOC can be applied to a task that depends on embodied motor control rather than high-order abstract cognition. To that end, we developed an experimental paradigm where we asked children to trace a series of predefined patterns on a tablet. Such a situation defies classical WOC studies because it is a complex motor task, which is difficult to parameterize or describe in simple terms, and whose errors are highly correlated (a deviation at any point in the line affects the future trajectory of the finger). It is also worth emphasizing that the task is intimately related to drawing, an important part of human culture and, with the appropriate experimental design and measuring tools, can be tested in naturalistic conditions beyond artificial laboratory settings. Although some studies have investigated collective problem-solving in tasks involving movement (24), our study is, to the best of our knowledge, the first one showing the implications of aggregating individual solutions to a sensory-motor task.

We present experimental results from hundreds of subjects tracing with their fingers a series of well-defined geometrical templates displayed on touchscreen tablets in a classroom setting. Using such “big behavioral data “(25) collected “outside the lab “(26), we examine the four main features that characterize the strong instances of WOC: (i) Whether individual trajectories of subjects can be aggregated to produce a more accurate description of the desired pattern, (ii) whether the improvement is a true “crowd “effect (requiring different individuals, as opposed to a single individual repeating the same task), (iii) whether the effect is strong enough so that the aggregate is more accurate than most of the individuals, and (iv) whether the effect is strong enough so that a crowd of low-skill individuals outperforms one high-skill individual. We find that all these conditions are met, providing the first evidence of Motor Wisdom of the Crowd.

## Results

### Collecting behavioral big data in a classroom setting

We asked 797 school students with ages between 6 and 18 years old, to trace with their finger several shapes using a custom-made drawing app (**Figure 1A**). After a few minutes of practice to familiarize themselves with the app, subjects were invited to reproduce five different geometric curves with varying levels of complexity, from ellipses to four-fold rose figures. A template of each shape was shown in the screen of the tablet (**Figure 1A**), and subjects were instructed to trace fluidly and continuously for 30 seconds, not excessively fast so as to avoid systematically overshooting the template but not excessively slow so as to avoid halts and brief jerky movements in trying to perfectly match the template. In other words, to simply produce good-enough tracing. All shapes were closed curves that could be traced repeatedly in a single stroke, and almost every subject traced each template several times during the 30 seconds of each experimental curve (**Figure 1B** and **Figure S1A**). Our experimental procedure allowed to test hundreds of children producing thousands of high-resolution trajectories (more than 24 hours of data) in naturalistic conditions (see **Methods** for more details).

**Figure 1.**
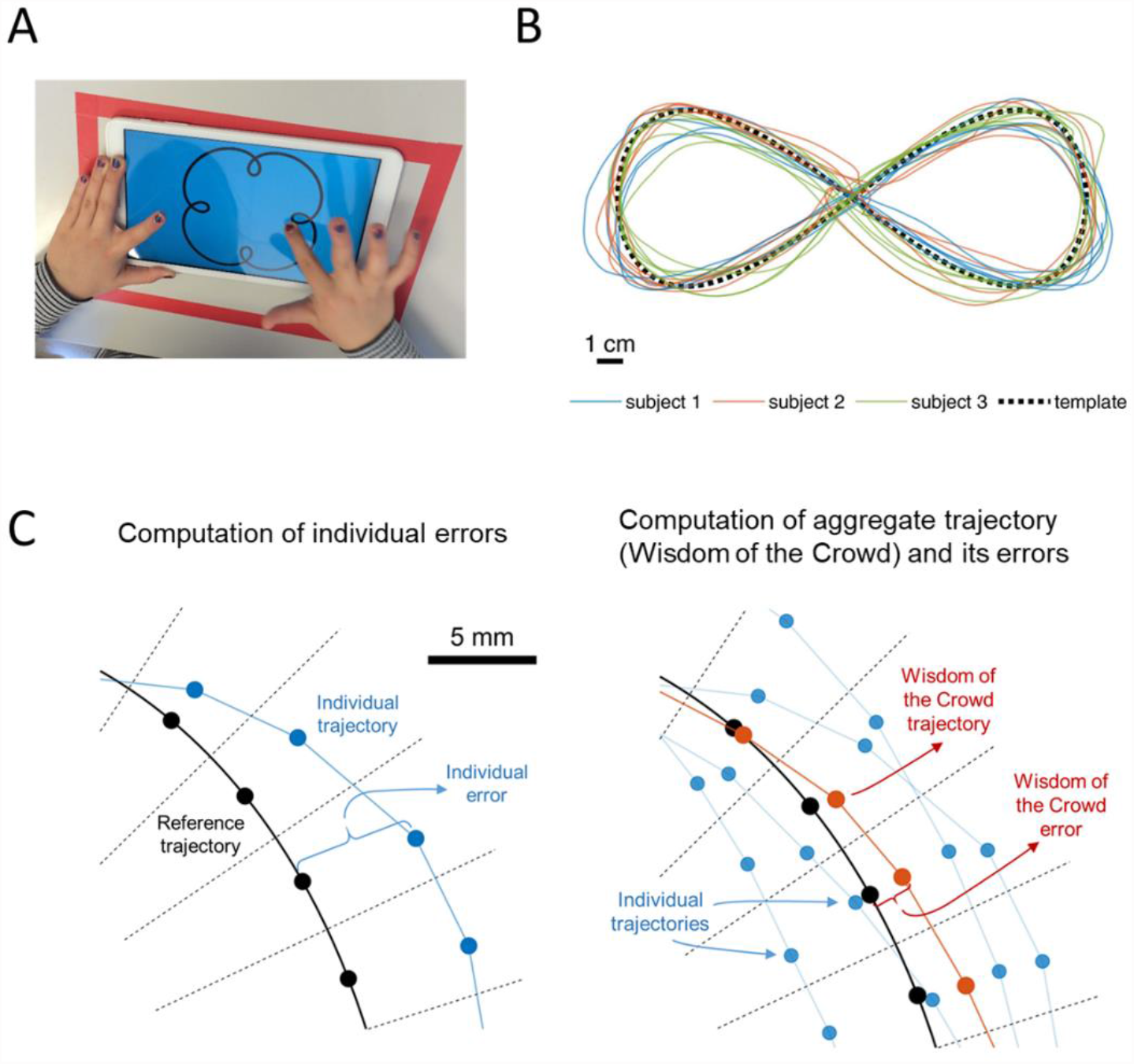
Description of the experiment and computation of errors. (**A**) One of the participants tracing on a tablet. The template was always present in thick black line on blue background. Participants were asked to trace the template with the index finger of their dominant hand continuously for 30 seconds. The procedure took place simultaneously in groups of 30 students in a classroom setting. (**B**) Example of template (black), and trajectories from three different subjects (blue, red, and green). (**C**) Method to compute errors and aggregate trajectories. Black lines represent the reference trajectory, which is divided in segments of equal length (limited by dashed lines). Each black dot is the median center of mass of the reference trajectory for each segment. Blue points: Experimental individual trajectories. The individual error is computed as the distance between the points for the individual trajectory and the respective centers of mass for the reference trajectory. Red points are the median center of mass of several individual trajectories (blue points) within the same segment. The Wisdom of the Crowd trajectory is formed with these averages (red line), and the error for the Wisdom of the Crowd is computed as the distance between these averages and the corresponding centers of mass of the reference trajectory. See **Figure S1** for a detailed description of trajectory subsampling, aggregation of individual trajectories, and computation of trajectory errors.

### Processing the trajectories to extract the Wisdom of the Crowd

In order to estimate the WOC of the trajectories traced by our subjects, the first step consisted in splicing the full trajectory of each subject into individual trajectories that represent a single pass over the template. This task turned out to be complicated for some complex shapes, and especially for low-accuracy trajectories that deviated very much from the template. To avoid biasing our results and increase the transparency of our analysis, we resorted to a simple approximate method: WeN determined that in most cases subjects traced each template at least 8 times, and divided the full trajectory of each subject into 8 segments of equal duration, which we then call “raw individual trajectories “(**Figure S1B**). While in most cases each raw individual trajectory contains more than a single pass over the template, this fact does not affect our conclusions and in fact strengthens them (see **Methods**).

Next, we subsampled the templates and processed the raw individual trajectories to facilitate their further analysis. We first took a set of reference points along each template (**Figure 1C** and **Figure S1C**). We then took the raw individual trajectory, assigned each of its points to the nearest reference point on the template, and found the median center of mass of each group of points (**Figure 1C** and **Figure S1D-H**). In this way, we obtained a subsampled individual trajectory (which we simply call “individual trajectory “), with one experimental point corresponding to each reference point of the template. From now on, we will refer to these subsampled individual trajectories simply as “individual trajectories “.

To compute WOC trajectories, we computed the median center of mass for the points associated to the same reference point from different individual trajectories (**Figure 1C, right** and **Figure S1I-K**).

To quantify the accuracy of a trajectory (either an individual trajectory or a WOC trajectory), we computed the distance between each of its points and the corresponding reference point of the template (**Figure 1C** and **Figure S1L**), obtaining the distance between them for each small region. Then, we used the average of these errors to quantify the overall error for the whole trajectory (**Figure S1M**).

We are now in position to establish whether the drawing task fulfills the four criteria of WOC mentioned above.

### Criterion i: WOC trajectories are more accurate than individual trajectories

We first investigated the accuracy of individual trajectories. Participants cared about properly tracing the templates but were not particularly motivated to be accurate, as we instructed them to trace quickly and fluidly without being too concerned about accuracy. Furthermore, our dataset includes data from very young children to late teenagers, whose motor skills are at different maturation stages. Consequently, individual trajectories showed a lot of dispersion (**Figure 2A**, blue).

**Figure 2:**
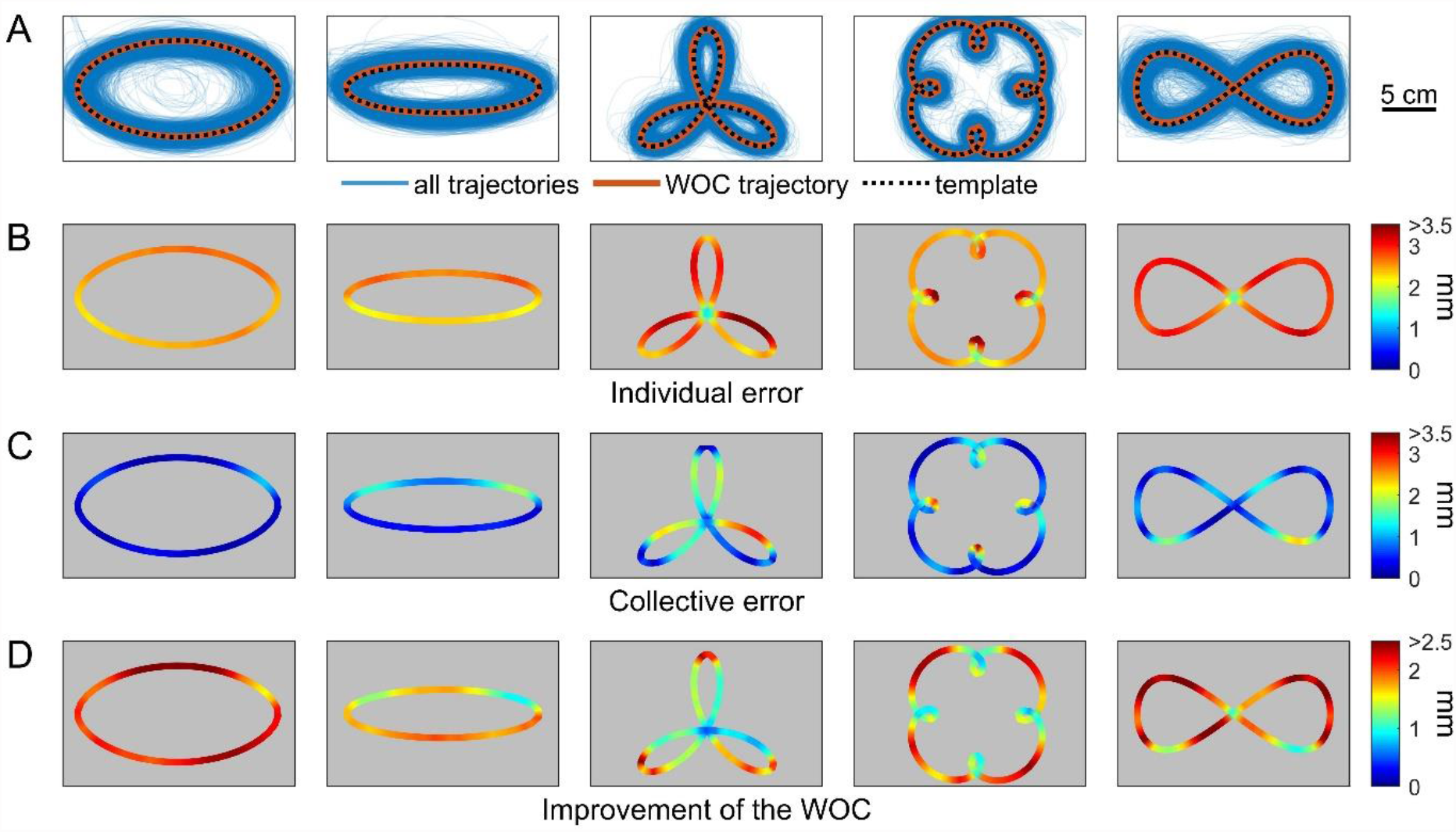
Wisdom of the Crowd in pattern tracing. (**A**) Original template (black dashed line), individual trajectories (blue), and WOC trajectory resulting from the aggregation of all individual trajectories (red). (**B**) Average error of individual trajectories for each region of the templates. (**C**) Error of the WOC trajectories for each region of the templates. (**D**) Improvement of the WOC over the individual trajectories for each region of the templates (i.e. average error of individual trajectories minus error of the WOC trajectory).

In spite of this inaccuracy at the individual level, WOC achieves remarkable accuracy. We built WOC trajectories for all shapes by taking one individual trajectory from each subject and aggregating all of them (**Figure 2A**, red). Average error (across all patterns) decreased more than twofold, from 2.72 mm for individual trajectories (**Figure 2B**) to 1.03 mm for WOC trajectories (**Figure 2C**). Improvement of WOC over individual trajectories was unequal over different portions of the templates (**Figure 2D**). Despite not showing a definite structure, one can notice interesting differences in the local improvement along different parts of the templates. In ellipses, major improvements take place in the straightest parts of the curve, where tracing speed is typically faster. While three-petal flowers show their WOC maximal improvement at the edges, four-petal flowers have it where curvature is minimal (rather than at the inner high-curvature turns). In both cases, however, the improvement corresponds to the most distal parts of the template taken globally. In the lemniscate, the WOC shows an intriguing global top-bottom asymmetry.

### Criterion ii: WOC accuracy improves with a diverse crowd

Aggregating several trajectories from a single subject should also lead to an improvement in accuracy, an effect termed “the crowd within “(27). Therefore, the improvement reported here for the WOC trajectories might not require a diverse crowd, but just be a consequence of aggregating multiple datasets (regardless of whether they come from the same subject or from different ones).

To elucidate whether our observation is a true effect of the crowd, we took advantage of the fact that each subject traced each pattern several times. While in the previous section we built our WOC trajectory from all of our subjects, here we studied how the error of the WOC trajectory changes as a function of how many individual trajectories are aggregated. We also compared the case of aggregating trajectories from the same individual (cycles while the same person draws the same pattern) or from different individuals. While in both cases the error decreases as we aggregate more trajectories, this decrease is much more rapid when the trajectories come from different individuals (**Figure 3A**), thus better supporting the WOC in movement trajectories.

**Figure 3:**
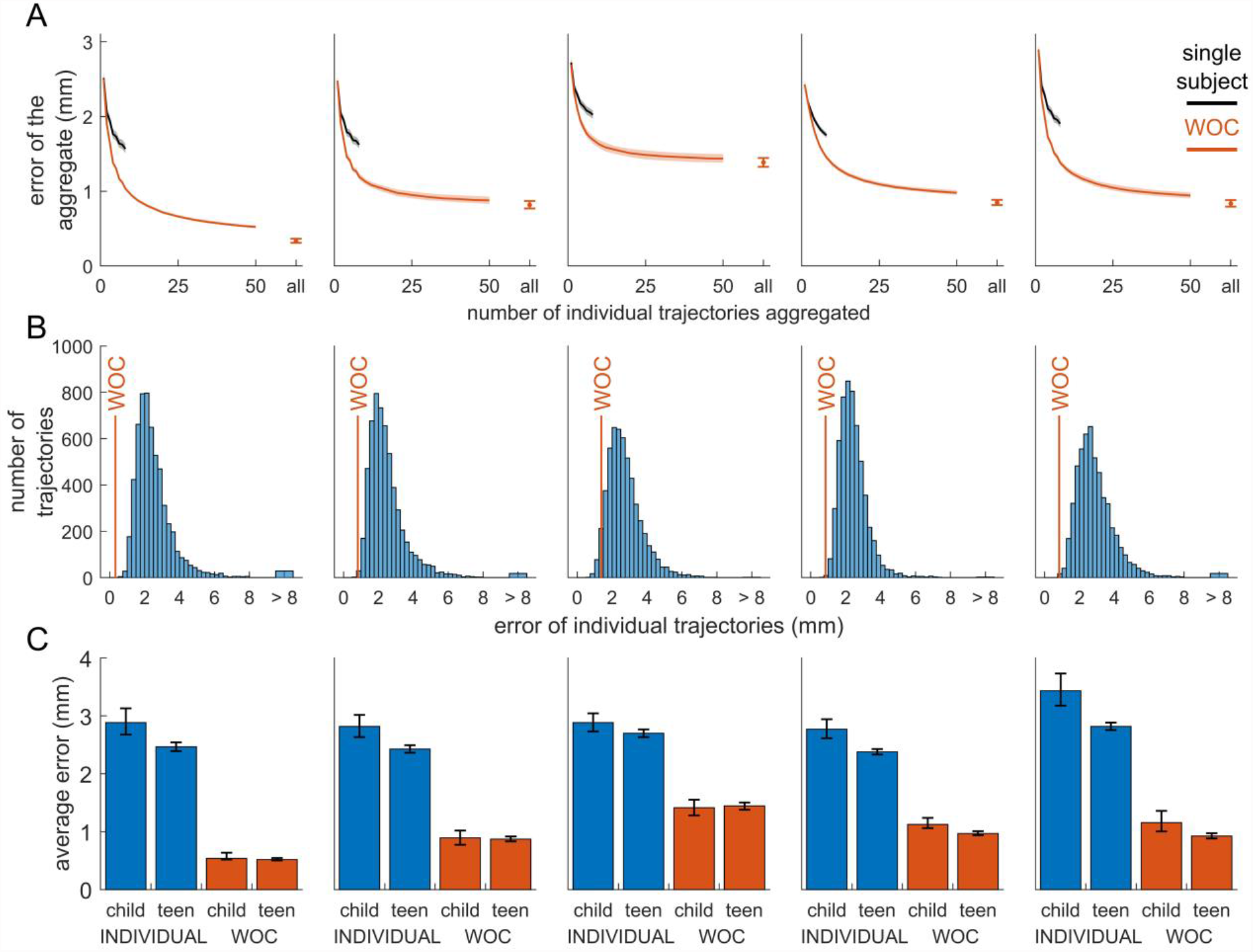
A strong case of Wisdom of the Crowd. Each column corresponds to one template, in the same order as shown in Figure 2. (**A**) Average error of the aggregate trajectory, as a function of the number of individual trajectories that are aggregated. Blue: Aggregated trajectories belong to the same subject. Red: Aggregated trajectories come from different subjects. Pale patches represent the 95% confidence interval (calculated via bootstrap). (**B**) Histogram of errors for individual trajectories, compared with the error of the WOC trajectory (red line). (**C**) Average individual error for children and teens (blue), and error of the WOC trajectory for children and teens (red). Error bars represent the 95% confidence interval (calculated via bootstrap).

This advantage of crowds over repeated trials of the same subject indicates that subjects tend to repeat their own errors. Different trajectories drawn by the same subject tend to deviate in the same regions and towards the same side, creating systematic biases. These biases are corrected when aggregating trajectories from different subjects, whose deviations are more balanced.

We also used this procedure to determine how many subjects are needed to achieve high accuracy. The error decreases monotonically as we add more trajectories from different subjects, dropping by almost half with only 10 subjects and starting to saturate after 50 subjects (**Figure 3A, red**).

### Criterion iii: WOC outperforms most individuals

A mere improvement of accuracy as we aggregate more trajectories would seem insufficient to support our claim of a WOC effect in motor control. In fact, it is a mathematical necessity that the error of an aggregate trajectory must be equal or smaller than the average error of the individual trajectories (14, 28).

What makes WOC such an important effect is the magnitude of the improvement. While there is no absolute threshold to consider that the effect is strong enough, a classical criterion is that the WOC estimate must be better than an overwhelming majority of the individual estimates. This was the case for example in Galton ‘s original demonstration of the effect, in which the arithmetic mean of individual guesses was better than every single individual estimate (29).

To investigate whether our WOC trajectories outperform the vast majority of individual estimates, we computed all individual errors for each template, and compared it with the error of the WOC estimate. In all cases the WOC estimate outperforms 99.5 % of the individual trajectories, except in one pattern in which it outperforms 95 % of the subjects (**Figure 3B**). Therefore, our dataset meets the criterion that the WOC estimate outperforms the vast majority of the individual estimates.

### Criterion iv: WOC of low-skill individuals beats the average high-skill individual

The last criterion of strong WOC, which is critical for its practical applicability, is that a crowd of low-skill individuals must outperform a single high-skill individual. To test this criterion, we first need to divide our population of subjects in groups with different expected skill levels. For tasks that require specific skills learned through education or professional training, high-skill subjects can be defined through their cultural level or profession. In the context of drawing, we considered testing professional painters and designers, but these professions usually have more to do with an aesthetic sense and the ability to master different drawing tools than with the motor control required to follow a predefined line with one ‘s finger.

However, our empirical approach allowed us to sample a large population of maturing subjects of different ages. Such a rich dataset offered a practical criterion to separate subjects by skill level: Age stratification. Our results include subjects of ages from 6 to 18 years (**Figure 4A**), and motor control develops during this period (especially between 6 and 10 years of age) (30–33). Indeed, our results show that individual performance improves with age (**Figure 4B**). Therefore, we defined low-skill individuals as young children (<10.5 years), and high-skill individuals as older children (>10.5 years) whose motor skills are comparably more developed (while this threshold maximizes the difference in individual performance between the two groups, our results hold regardless of the threshold chosen, **Figure S2**).

**Figure 4:**
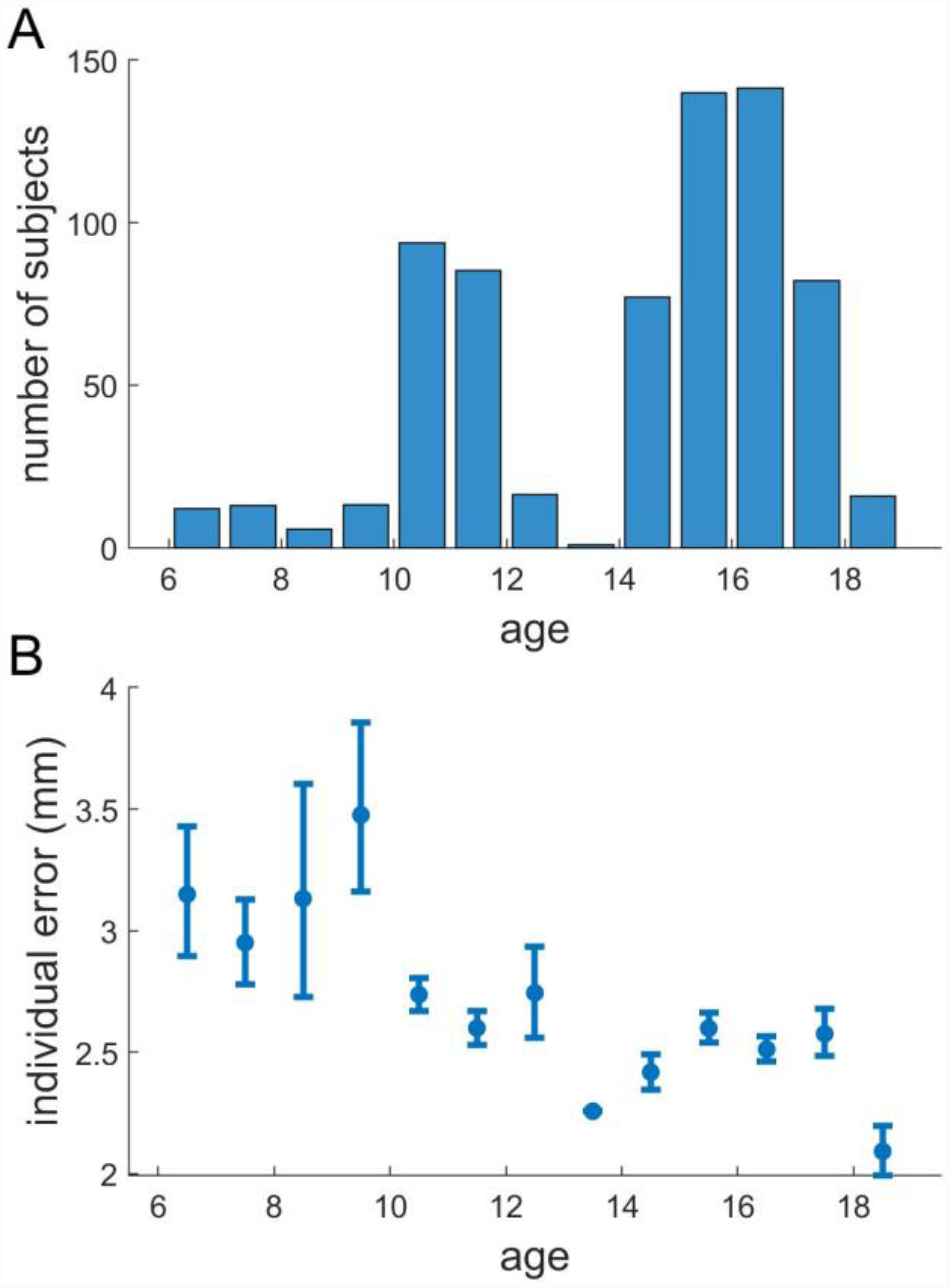
Effect of age. **(A)** Number of subjects of each age in our dataset. **(B)** Average error of the individual trajectories (averaged over all segments of each template, over individuals with same age, and over the 5 templates), as a function of age. Error bars represent the 95% confidence interval (calculated via bootstrap).

As a confirmation to this criterion, individual estimates are better for older children than for younger ones (**Figure 3C**, compare bars 1 and 2 of every template). The key question is then whether a crowd of young children outperforms the average older child. Remarkably, we found this to be true for every template (**Figure 3C**, compare bars 2 and 3 of every template). Therefore, our dataset meets the last criterion of the strongest versions of WOC: A crowd of low-skill individuals outperforms the average high-skill individual.

### High-skill individuals play little role in the Motor Wisdom of the Crowd

We also found an interesting and unexpected result. When comparing the results from WOC estimates from old and young children, we found almost no difference. Only for 2 of the 5 templates the WOC estimates are better for older children than for younger ones (**Figure 3C**, compare bars 3 and 4 of every template). This result indicates that, when recruiting a crowd to perform a WOC estimate, there is no benefit in selecting high-skill individuals (34). In other words, one does not need to include (motor) “experts “to achieve the (motor) Wisdom of the Crowd.

We also asked the question of whether different criteria to define high-skill or low-skill individuals could give different results. To answer this question, we simulated criteria that estimated individual skill with different accuracies, from those capable of selecting the very best individual, to benchmarks incapable of distinguishing skill. Our results turned out to be very robust: For a high-skill individual to outperform the crowd, the selection method must be able to identify accurately the top 1.73% performers (**Figure S3**).

### The method to aggregate trajectories has little impact on the results

An important practical question is how to aggregate the individual responses to produce the WOC estimate (7, 14), and in particular whether to use the mean or the median, considering that the latter can counteract the effect of outliers (35, 36). All our previous analyses use the median to compute centers of mass, as we detected some subjects with very large deviations from the templates. We re-did our analysis using the mean instead of the median and compared both methods, to find very little difference between them (**Figure S4**). This result indicates that, even though some subjects are clear outliers, the distribution of trajectories around the true pattern has relatively little skew (**Figure 2A**, note how the density of traces is nearly symmetric along the templates).

## Discussion

Here we tested whether Wisdom of the Crowd can take place in a context far removed from those studied so far. By asking our subjects to trace a complex trajectory rather than estimating a number, we investigated a procedure that does not consist in making an explicit rational estimate but in performing an implicit embodied motor task. We tested hundreds of children drawing in a custom-made tablet app, collecting a large amount of precise and quantitative data to test the effect of aggregating independent individuals for a better collective performance.

We found that tracing geometrical patterns manifests all the characteristics of Wisdom of the Crowd: (i) accuracy improves when aggregating several trajectories, (ii) these trajectories must come from different subjects, (iii) the aggregate trajectory outperforms most individual ones, and (iv) a crowd of low-skill individuals outperforms the typical high-skill individual. Our results thus extend the concept of Wisdom of the Crowd from its classical application of quantity estimation to the realm of motor control and embodied cognition.

Our findings suggest that Wisdom of the Crowd may be applicable with a greater generality than previously thought, because they address outstanding theoretical concerns: For the WOC estimate to be accurate, the individual estimates must follow a probability distribution whose average (either the mean or the median) matches the true value. A general concern was that the tasks typically chosen, such as number estimation, might share some characteristics that made them fulfil this requirement, which would not be met when trying to extend Wisdom of the Crowd to other contexts. Here we have tested Wisdom of the Crowd on a completely different context, in which the object to be estimated is not a number but a complex trajectory, and the estimation procedure is not a rational thought process but a motor task. Our results indicate that the conditions required for Wisdom of the Crowd are met with great generality, and call for a more systematic exploration across tasks of different nature.

The high accuracy of the motor Wisdom of the Crowd is surprising. Given the complex nature of the task, the extreme inaccuracy of some trajectories, and the fact that errors accumulate (a deviation at one point in the trajectory affects its future), we did not expect that simple aggregation of trajectories would result in such accurate estimations. This high accuracy opens the door to applying similar aggregation methods to other tasks with similar characteristics, such as finding the optimal trajectory of a vehicle (37) or in the context of skill improvement in amateur and professional athletes.

Our results suggest a potential application in the study of human symbols. Letters, numbers and other symbols are written differently by different people (38), and their shape changes significantly over time and from region to region (39). These changes are well documented qualitatively, and recent techniques have been developed to characterize them quantitatively (40, 41). These techniques are usually based on aggregating images, regardless of the trajectories that the writers followed to trace each symbol. Our results indicate that it would be possible to compute an average trajectory for a given symbol across a population. The methodology would need to be different to the one presented in this paper, with trajectory estimation and alignment posing important methodological challenges. But the surprisingly high accuracy found in our dataset indicates that aggregating symbols written by different people may provide an intelligible average tracing trajectory, which would facilitate the study of how handwriting changes in space and time.

## Methods

### Hardware

Thirty Android touch-screen tablets were used for the behavioral experiments. The tablet brand was Samsung Galaxy Tab A6 (size: 254 × 164 × 8mm, Android version 8.1.0 and API level 26). Prize per tablet was less than 200$. The display has a 10.1 “PLS LCD screen, with dimensions 216 × 135mm, and a resolution of 1920 × 1200px. The tablet has a capacitive touch-screen, and registers touch by a finger or a capacitive stylus, with resolution equal to the display resolution. Maximum screen refresh rate is 60Hz, and maximum sampling rate of touch events is close to 85Hz.

### Software

The app was programmed in Android Studio (version 3.3.2) in the Kotlin programming language (26), and tailored specifically for accuracy, efficiency and robustness in out-of-the-lab experiments with children. It can be freely downloaded (https://github.com/adam-matic/KinematicCognition), and also edited for different experimental purposes.

### Experimental procedures

A total of 851 subjects participated in the experiment, most of which were school students between 6 and 18 years old (56% female, 10% left-handed, see **Table S1** for number of adults). All experiments were performed during the 2019 Brain Awareness Week (from March 11 to 15, 2019). Students arrived in groups of about 30 individuals, belonging to the same school class. Classes belonged to several different schools in the area of Alicante, Spain. There were no specific selection criteria for schools and classes beyond their willingness to participate in our experiment and a more or less homogeneous sampling of ages and locations. Groups were assigned different time slots throughout the morning. The experiments took place in a regular small classroom (with a capacity for 30 students) in a building adjacent to the Instituto de Neurociencias de Alicante, Spain. We used 30 tablets (a single tablet per children), placed on the tables in the classroom. Each experimental session had an overall duration of about 15 minutes.

Students were invited to enter the classroom and sit down wherever they wished. Then, before starting the experiments, students were greeted and briefly told about the overall goal of the study (they already knew some details because they had an explanation of the activity days before by their teachers at their own local schools and, also, their parents had been previously informed and asked for written consent to record their children finger trajectories). The explanation was generic, namely, a brief and fun experiment was going to take place where they would simply need to draw and trace specific geometric figures in order as they would appear on the tablet screen, helping scientists, to study motor control (this fitted well in the Brain Awareness Week, since each group also received a related outreach talk, so that they could not only listen about scientific experiments but actually also participate in them).

Students were requested to use the index finger of their dominant hand to draw and trace on the tablet screen (rather than using tablet pens). They were also asked to avoid touching the screen with the other hand, or to move the tablet from where it was placed when they entered the room. Before starting the experiment, they all tapped on the screen at the custom-made app logo of the experiment.

First, a simple screen opened where they were asked whether their dominant hand was left or right, their age by scrolling on the date of birth, and their gender. After the information is complete, the app allows two options: “Practice “or “Experiment “.

Second, the participants did a trial exercise before the actual experiment, where similar curves appeared as to the ones they would encounter later. This part was key for them to familiarize with the tasks they would need to accomplish next. In particular, they were instructed not to separate the finger from the screen until each task was over, and to perform fluid movements, avoiding delineating too slow or too fast. They could at this point ask questions, before the experiment took place.

Third, after such trials, oral instruction prompted the students to start all the “Experiment “part at the same time. The experimental part consisted of a series of exercises, or tasks, all automatically concatenated with brief pauses in between. In this way, we avoided having to verbally interrupt the whole classroom with various unnecessary (and potentially distracting and confusing) instructions (specially for the younger children) to start the many different drawing and tracing tasks. Moreover, although sitting next to each other, the performance of each participant was purely individual, since every student was concentrated in their own tablet, not paying attention to their neighbor ‘s.

Three different classes of tasks were presented to the participants: tracing, tracking and scribbling. Every class comprised different exercises, each one with a duration of 30 seconds, with a 7-second pause in between. Thus, the whole motor control experience was brief, avoiding distractions or loss of interest by the children. A visual summary of the experimental dataset can be found here: https://youtu.be/rz-TWk_6HSU.

In this study we only analyzed the first class of task (tracing), where participants had to delineate with their finger a black curve on a blue screen. The curves shown were an oval (or ellipse), a thinner oval (larger eccentricity), three-petal and four petal-flowers (both based on Huh ‘s pure frequency curves (42)), and an infinite symbol (lemniscate).

When the experiment finished, students were thanked again, and invited to leave the room to continue enjoying the Brain Awareness Week parade at the Instituto de Neurociencias. Experimental procedures were approved by the Institutional Review Board (under project registration number 2019.111.E.OIR) and followed the required guidelines on participation and personal data protection.

### Data cleaning

We removed anomalous data following three criteria: (1) Trajectories that lasted less than 25 seconds. (2) Trajectories whose standard deviation (in either the horizontal or vertical dimension) was less than half the standard deviation of the points of the template. (3) Trajectories with jumps between two consecutively recorded points greater than 1/4 of the dimensions of the touchscreen, which in most cases were due to a malfunctioning by the tablet or the subject touching the screen with both hands simultaneously. See **Table S1** for the number of full trajectories originally stored, the number of trajectories that did not meet each of the filtering criteria (some trajectories did not meet more than one criterion), and the number of trajectories that were finally used for the analysis. In sum, a total of 3485 trajectories were analyzed, each with a duration longer than 25 seconds, which yields an estimate of a total of more than 24 hours of high-resolution quantitative measurements of human drawing in naturalistic conditions.

### Definition of individual trajectories

Each subject traced the template repeatedly during the 30 seconds allotted for the task, so each trajectory contains several passes over the template. The ideal definition for “individual trajectory “would be a single pass over the template, but finding the exact point in which the trajectory finishes one pass and starts the next is problematic: When the trajectory is a very poor approximation to the template, it is unclear when we should consider that an individual trajectory has ended. Therefore, attempts to divide the trajectories in this way would either force us to discard the less accurate trajectories, or could result in systematic biases affecting differently the high- and low-accuracy trajectories.

For this reason, we chose a simple definition of individual trajectory: We divided each trajectory in 8 segments of equal duration, and we took each of these segments as an individual trajectory. While this is only an approximation, it has the advantage of being a simple and transparent method and avoiding biases among trajectories of different accuracy.

Because of this approximate definition, in many cases an individual trajectory contains only part of the template or more than one pass over some parts of the template. In most cases subjects completed more than 8 full passes over the whole template (even for the most complex shapes) so on average our individual trajectories contain more than one pass. We made this choice to be conservative: Since our individual trajectories in fact contain more than one pass over each template (on average), the individual errors we report are underestimated (a more accurate definition of individual trajectories would remove the aggregation that is taking place in the regions with more than one pass, which reduced the error). Therefore, the effect of Wisdom of the Crowd in our dataset is, if anything, underestimated.

### Subsampling of individual trajectories

Raw individual trajectories contained between 200 and 215 points. Before any analysis, we sub-sampled the templates into a set of reference points (50 reference points for the two ellipses and 100 reference points for the rest). Then, we sub-sampled the raw individual trajectories to ensure that each point of a subsampled individual trajectory corresponded to one reference point of its corresponding template. We did this by finding the median center of mass of all points of the raw individual trajectory nearest to each reference point of the template (similar results are obtained by using the arithmetic mean instead of the median, as shown in **Figure S4**). See **Figure S1** for a step-by-step description of this process. The term “individual trajectory “always refers to the sub-sampled one (we use “raw individual trajectory “to name the original trajectory before sub-sampling).

### Aggregation of individual trajectories

To aggregate several individual trajectories (regardless of whether they belonged to the same subject or to different subjects), we computed the median center of mass of the points of each individual trajectory corresponding to the same reference point in the template (**Figure 1C**). The final result is an aggregate trajectory that has the same number of points as the number of reference points of the template (see **Figure S1**).

### Computation of errors

To compute the error of any trajectory (either an individual trajectory or an aggregated one), we found the Euclidean distance between each of its points and the corresponding reference point of the template (**Figure 1C** and **Figure S1**). These distances are used to represent the colors in **Figure 2B-D**, showing the average error for each region of each template across all the trajectories. The total error of a trajectory is computed as the arithmetic mean of all the distances to the reference points.

To compute the average error when aggregating individual trajectories from the same subject (**Figure 2E**, blue lines), we created all possible sets with a given number of individual trajectories from each subject. For example, when aggregating 3 individual trajectories, for each subject there are 56 different combinations out of the total of 8 individual trajectories, so there are 56 different sets of 3 individual trajectories per subject. The error for the template is determined as the average of the errors over all sets of each subject, and then over all subjects.

To compute the average error when aggregating individual trajectories from different subjects (as for example in **Figure 2E**, red lines), it would not be possible to compute all possible combinations (there would be too many). Therefore, we performed a random sample: We randomly drew the desired number of subjects from the database, and for each subject we randomly chose one of the individual trajectories. Then, we aggregated the individual trajectories, and computed the error of the aggregate one. We repeated this process 10000 times, and computed the arithmetic mean of all errors.

To compute the average error across templates in **Figure 3**, we first computed the expected error for each of the 5 templates, and then computed the arithmetic mean of these 5 error values.

### Computation of confidence intervals

To compute the confidence intervals for the errors of individual and aggregated trajectories (**Figures 2E and 2G**), we used bootstrap (43). First, we created “virtual experiments “by randomly drawing subjects with repetition until we reach the total number of subjects. Therefore, each virtual experiment consisted of the same number of subjects, but due to the random sampling some of our original subjects may be missing, and some subjects may be present more than once. Then, we recreated all our analysis on each of these virtual experiments. We repeated this full process with 500 virtual experiments, and our confidence intervals represent the region that contain 95% of these results.

## Contributions

AGM conceived the experiments. RZL, AM and AGM designed the experiments. RZL performed the experiments. AGM and AM designed the tablet app for big behavioral data collection in the real world. AM programmed the tablet app and processed the trajectory data. GM, APE designed the Wisdom of the Crowd analysis. GM performed the analysis and made the figures. GM, APE and AGM interpreted the results. GM and APE wrote the first full draft of the manuscript. All authors contributed to the writing of the final version of the manuscript.

## Acknowledgements

We thank María del Carmen Lillo Navarro for assistance during some of the experiments. We are also grateful to Alicia Ferri and Diego Echevarría for the general organization of the Brain Awareness Week at the Instituto de Neurociencias de Alicante. We thank members of IVEP at the Research Center for Animal Cognition in Toulouse, France, for feedback on a previous version of the manuscript.

