## Supplementary Figures and Table for "The Motor Wisdom of the Crowd"

### Figure S1: Analysis of Trajectories

| | |
|---|---|
| A 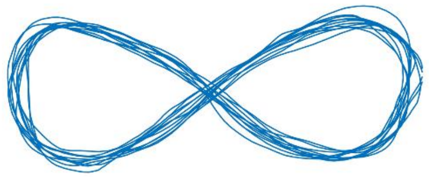 | Each subject traced the template several times over the duration of the task.<br><br>**Blue line:** Full trajectory for one subject |
| B 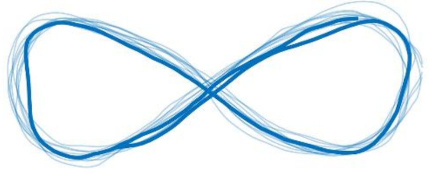 | Each full trajectory is divided into 8 segments of equal duration, and each of these segments is called "raw individual trajectory".<br><br>**Thin line:** Full trajectory for one subject.<br>**Thick line:** One individual trajectory, covering 1/8 of the full trajectory. |

| | |
|---|---|
| C<br>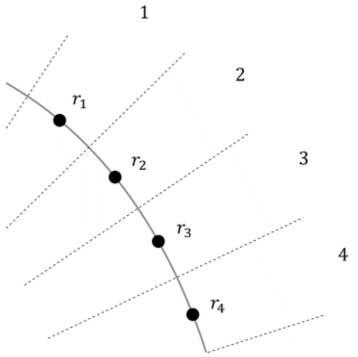<br> | We chose a set $\{r_i\}_{i=1}^{n_r}$ of $n_r$ equi-spaced reference points along the template.<br><br>**Black line:** Template<br>**Black points:** Reference points, $\{r_i\}_{i=1}^{n_r}$<br>**Dashed lines:** Limits between the regions of space closest to each reference point. |
| D<br>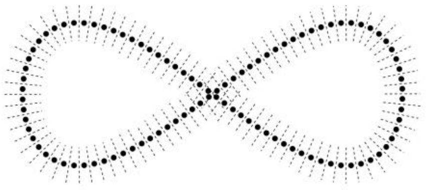 | We have $n_r$ = 50 points in the case of the two ellipses, and $n_r$ = 100 points for the other templates (which were geometrically more complex and included crossings).<br><br>**Black points:** Reference points, $\{r_i\}_{i=1}^{n_r}$<br>**Dashed lines:** Limits between the regions of space closest to each reference point. |
| E<br>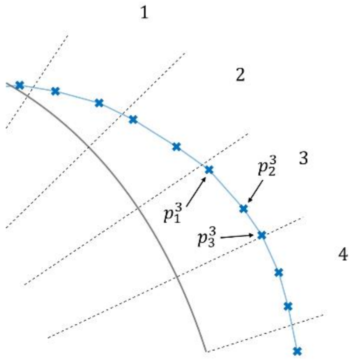<br> | For each raw individual trajectory we found all points $\{p_k^i\}_{k=1}^{n_i}$ closest to each reference point $r_i$ of the template.<br><br>**Black line:** Template<br>**Dashed lines:** Limits between the regions of space closest to each reference point of the template.<br>**Blue crosses:** Experimental points, $\{p_k^i\}_{k=1}^{n_i}$. The three points marked by the arrows are the ones closest to reference point number 3. |

F

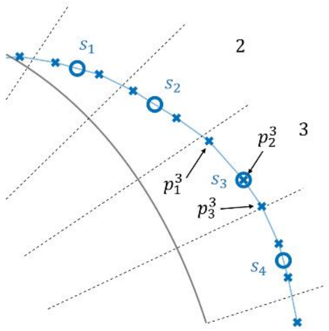


We found the median center of mass for each set of experimental points closest to one reference point. The median center of mass is the point whose coordinates are the median of the coordinates of the original points: $s_i = median(\{p_k^i\}_{k=1}^{n_i})$.

**Blue circles:** Median center of mass for the experimental points in each region ($s_i$).

G

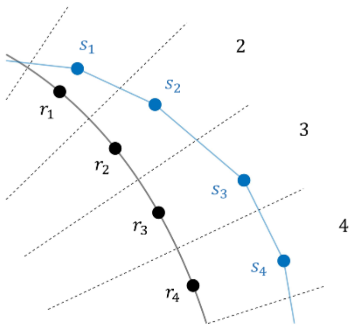


This method provided a subsampled individual trajectory $\{s_i\}_{i=1}^{n_r}$, each of whose points $s_i$ corresponds to one reference point $r_i$ in the template.

H

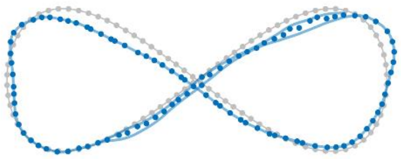

We always performed this subsampling in the first place to ensure that each point of an individual trajectory corresponded to one reference point of the template. All references to "individual trajectories" refer to these sub-sampled individual trajectories (whenever an individual trajectory is not subsampled, we explicitly refer to it as "raw individual trajectory").

Note that if our individual trajectory contains more than one pass over the same region, our method will also aggregate the different passes, taking advantage of the "crowd within" effect.

**Gray:** Template
**Blue:** Individual trajectory (after subsampling)

I

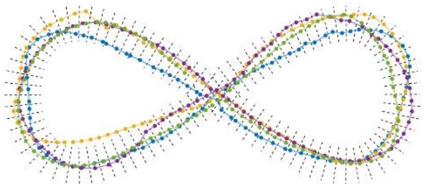

We now turn to how we aggregate several individual trajectories (either from the same subject or from different subjects).

**Dashed lines:** Regions closest to each reference point of the template
**Colored points and lines:** Four different individual trajectories.

J

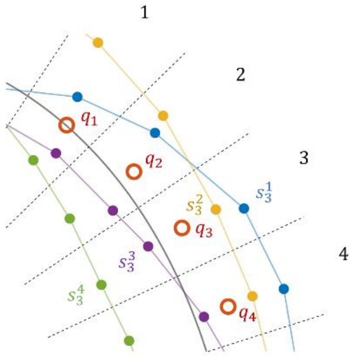


We computed the aggregate trajectory $\{q_i\}_{i=1}^{n_r}$ by finding the median center of mass of the point of each individual trajectory corresponding to the same reference point in the template: $q_i = median(\{s_i^j\}_{j=1}^{m_s})$, with $m_s$ the number of individual trajectories aggregated.

**Dashed lines:** Regions closest to each reference point of the template
**Colored points and lines:** Four different individual trajectories.
**Red circles:** Points of the aggregate trajectory

K

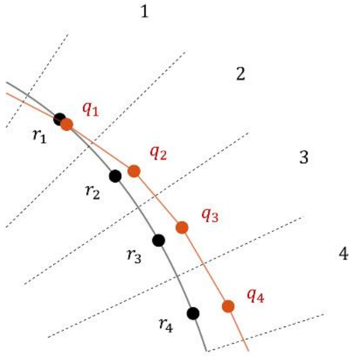


The final result is an aggregate trajectory that has the same number of points as the number of reference points of the template ($n_r$).

<table>
<tr>
<td>L<br>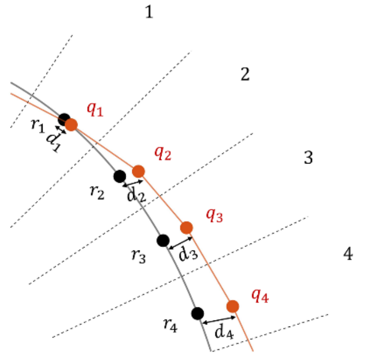<br></td>
<td>To compute the error of any trajectory (either an individual trajectory or an aggregated one), we found the Euclidean distance between each of its points $s_j$ and the corresponding reference point $r_i$ of the template: $d_i = |r_i - s_i| = [(x_i^r - x_i^s)^2 + (y_i^r - y_i^s)^2]^{1/2}$.</td>
</tr>
<tr>
<td>M<br>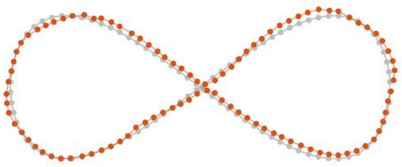</td>
<td>The total error of a trajectory is computed as the arithmetic mean of all the distances to the reference points: $\varepsilon = (1/n_r)\sum_{i=1}^{n_r} d_i$.</td>
</tr>
</table>

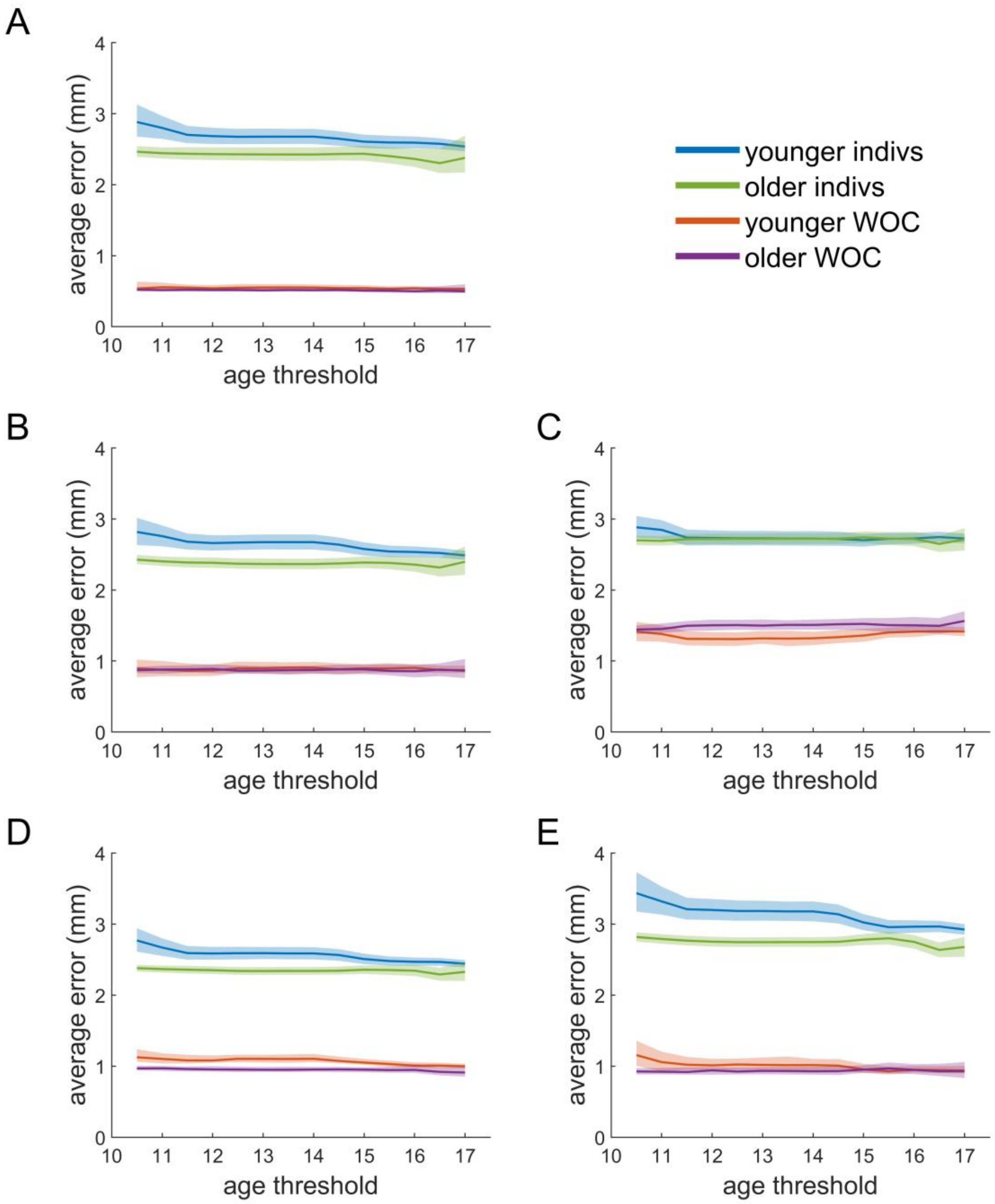


**Figure S2. Error of subjects younger or older than a threshold.** Average error of the individual trajectories for all subjects younger (blue) and older (green) than the age threshold indicated in the x-axis. Shown also, the error when aggregating one trajectory from each of the subjects younger (red) and older (magenta) than the age threshold. Pale patches represent the 95% confidence interval (calculated via bootstrap). The templates are in the same order as shown in **Figure 2**: **(A)** Ellipse; **(B)** Long ellipse; **(C)** Three-petal flower; **(D)** Four-petal flower; **(E)** Lemniscate.

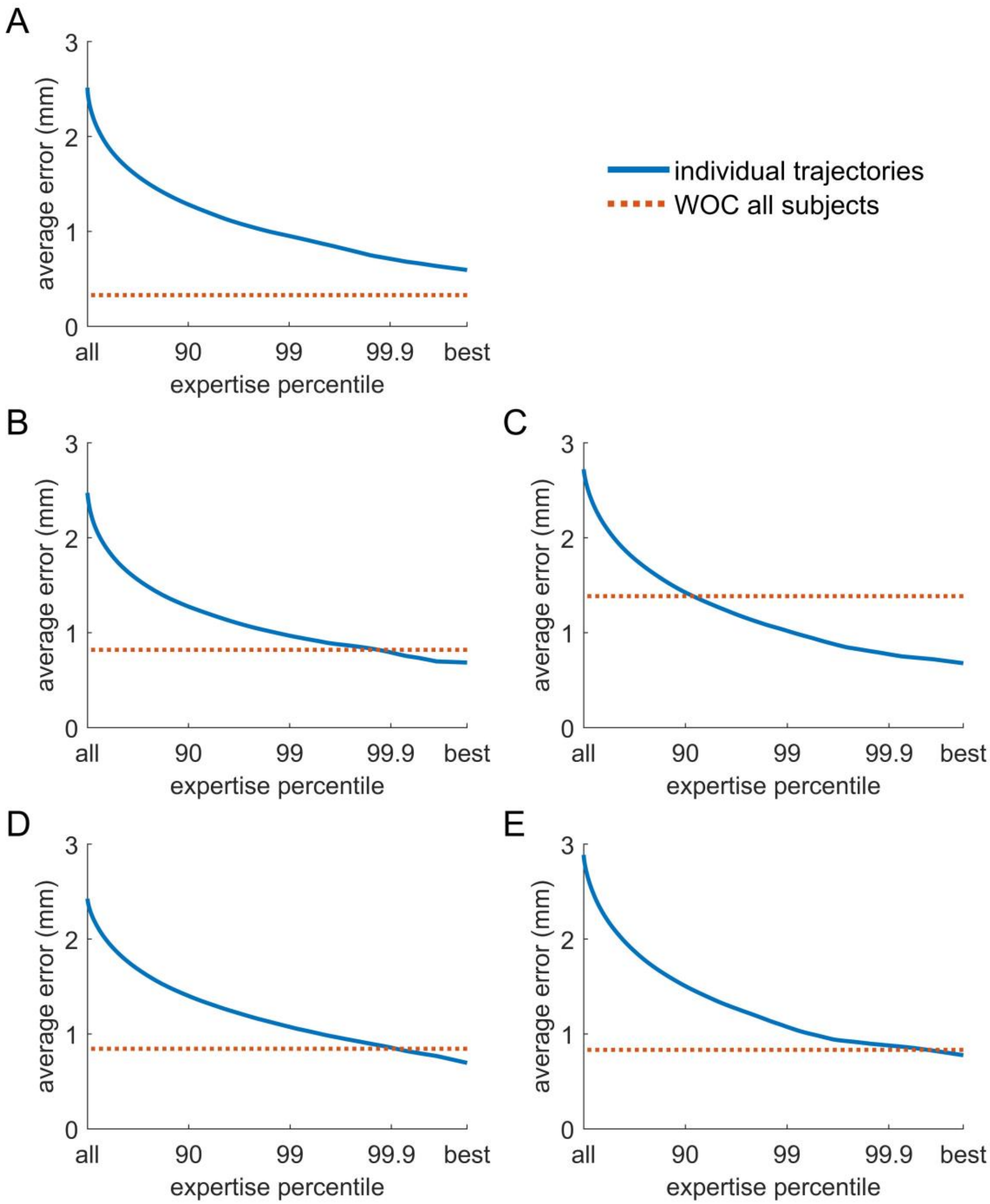


**Figure S3. Error when selecting the best performing subjects.** First, all individual trajectories are sorted by their error in descending order, so each one can be assigned to an expertise percentile. Then, for each of the percentiles indicated in the x-axes, the trajectories with an error equal or lower than that of the percentile are selected. Blue: Average error of individual trajectories located at the expertise percentile or higher. Red: Average error when aggregating one individual trajectory from each subject in the dataset, computed as for **Figure 3A**. The templates are in the same order as shown in **Figure 2**: **(A)** Ellipse; **(B)** Long ellipse; **(C)** Three-petal flower; **(D)** Four-petal flower; **(E)** Lemniscate.

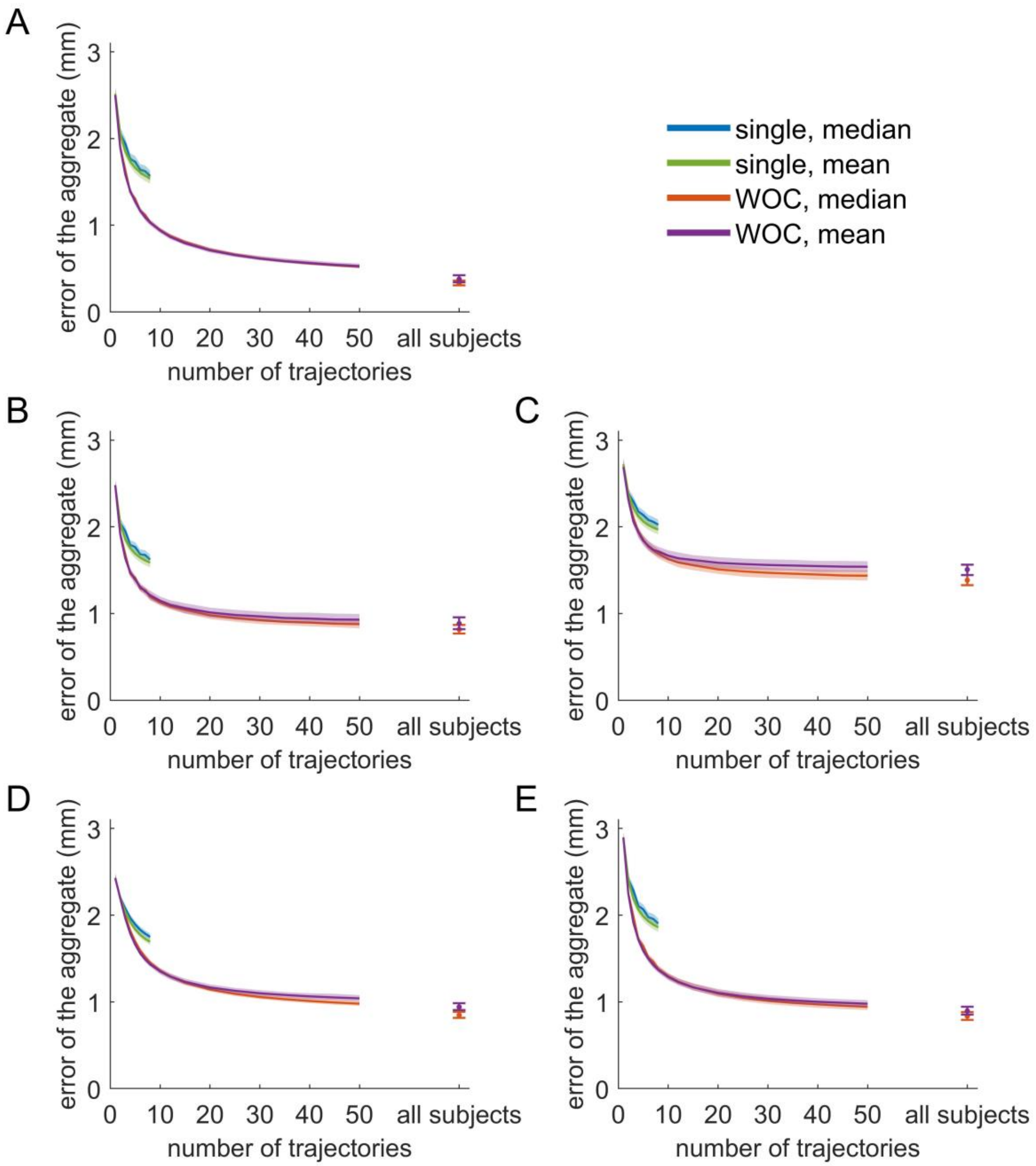


**Figure S4. Using the arithmetic mean instead of the median to aggregate trajectories.** Blue: Aggregation using the median of trajectories belonging to the same subject. Red: Aggregation using the median of trajectories that come from different subjects. Green: Aggregation using the arithmetic mean of trajectories belonging to the same subject. Magenta: Aggregation using the arithmetic mean of trajectories that come from different subjects. Pale patches represent the 95% confidence interval (calculated via bootstrap). Error of the aggregate trajectories as a function of the number of individual trajectories that are aggregated, computed as for **Figure 3A**. The templates are in the same order as shown in **Figure 2**: **(A)** Ellipse; **(B)** Long ellipse; **(C)** Three-petal flower; **(D)** Four-petal flower; **(E)** Lemniscate.

| Template | Original number | Older than 18 | Too short | Too narrow | Big jumps | Final number |
|---|---|---|---|---|---|---|
| **Ellipse** | 851 | 54 | 46 | 19 | 38 | 728 |
| **Flat Ellipse** | 810 | 50 | 14 | 1 | 40 | 711 |
| **Flower 3** | 816 | 50 | 32 | 5 | 77 | 670 |
| **Flower 4** | 810 | 49 | 27 | 7 | 45 | 700 |
| **Lemniscate** | 810 | 49 | 21 | 3 | 73 | 676 |

**Table S1. Rejected trajectories.** For each template (first column), shown is the original number of trajectories stored (second column), how many trajectories did not meet each filtering criterion (third to last but one columns), and the total number of trajectories that remained and were therefore used for the analysis (last column).
